# Disentangling sources of selection on exonic transcriptional enhancers

**DOI:** 10.1101/024000

**Authors:** Rachel M. Agoglia, Hunter B. Fraser

## Abstract

In addition to coding for proteins, exons can also impact transcription by encoding regulatory elements such as enhancers. It has been debated whether such features confer heightened selective constraint, or evolve neutrally. We have addressed this question by developing a new approach to disentangle the sources of selection acting on exonic enhancers, in which we model the evolutionary rates of every possible substitution as a function of their effects on both protein sequence and enhancer activity. In three exonic enhancers, we found no significant association between evolutionary rates and effects on enhancer activity. This suggests that despite having biochemical activity, these exonic enhancers have no detectable selective constraint, and thus are unlikely to play a major role in protein evolution.

---

Beyond their essential role in specifying amino acid sequences, exons can play additional roles in the regulation of transcription, translation, splicing, and mRNA stability. For example, transcriptional enhancers located within exons can regulate the expression of their respective genes, or neighboring genes, in a tissue-specific manner (Birnbaum et al. 2012; Ritter et al. 2012).

It is plausible that exonic enhancers may be subject to purifying selection to preserve their function, leading to greater conservation than in exons without regulatory roles (Lin et al. 2011; Suzuki & Saitou 2011). An initial investigation of conservation at exonic enhancer sites (Stergachis *et al.* 2013) examined the conservation of fourfold degenerate bases, both inside and outside exonic transcription factor (TF) binding sites. This analysis found that exonic enhancers are more conserved than non-regulatory exons, and the authors concluded that exonic enhancers have a strong influence on the trajectory of protein evolution. Recently however, a reevaluation of this claim argues that when using a different metric to measure conservation, there is no detectable difference in conservation between TF binding and non-binding sites within exons (Xing and He 2015). Both of these contradictory studies focused on genome-wide trends, and thus may be susceptible to biases caused by confounding factors that were not taken into account. Indeed, the latter study claims that the original finding of conservation was primarily due to higher GC content in exonic TF binding sites. Whether additional confounders may exist is still an open question.

We decided to investigate this question from a different perspective, by developing a method for comparing the relative roles of protein sequence and enhancer function in selection acting on exonic enhancers (Fig 1). Building on the method introduced by Smith et al. (2012) for detecting selection on regulatory elements, we tested whether the substitutions that have occurred during the evolution of specific exonic enhancers are best explained by selection on protein sequence, enhancer activity, or a combination of both. This is possible when, for each possible single-nucleotide variant (SNV), we have experimental data measuring its effect on enhancer activity. More formally, we fit a linear model of the form:

**Fig. 1.**
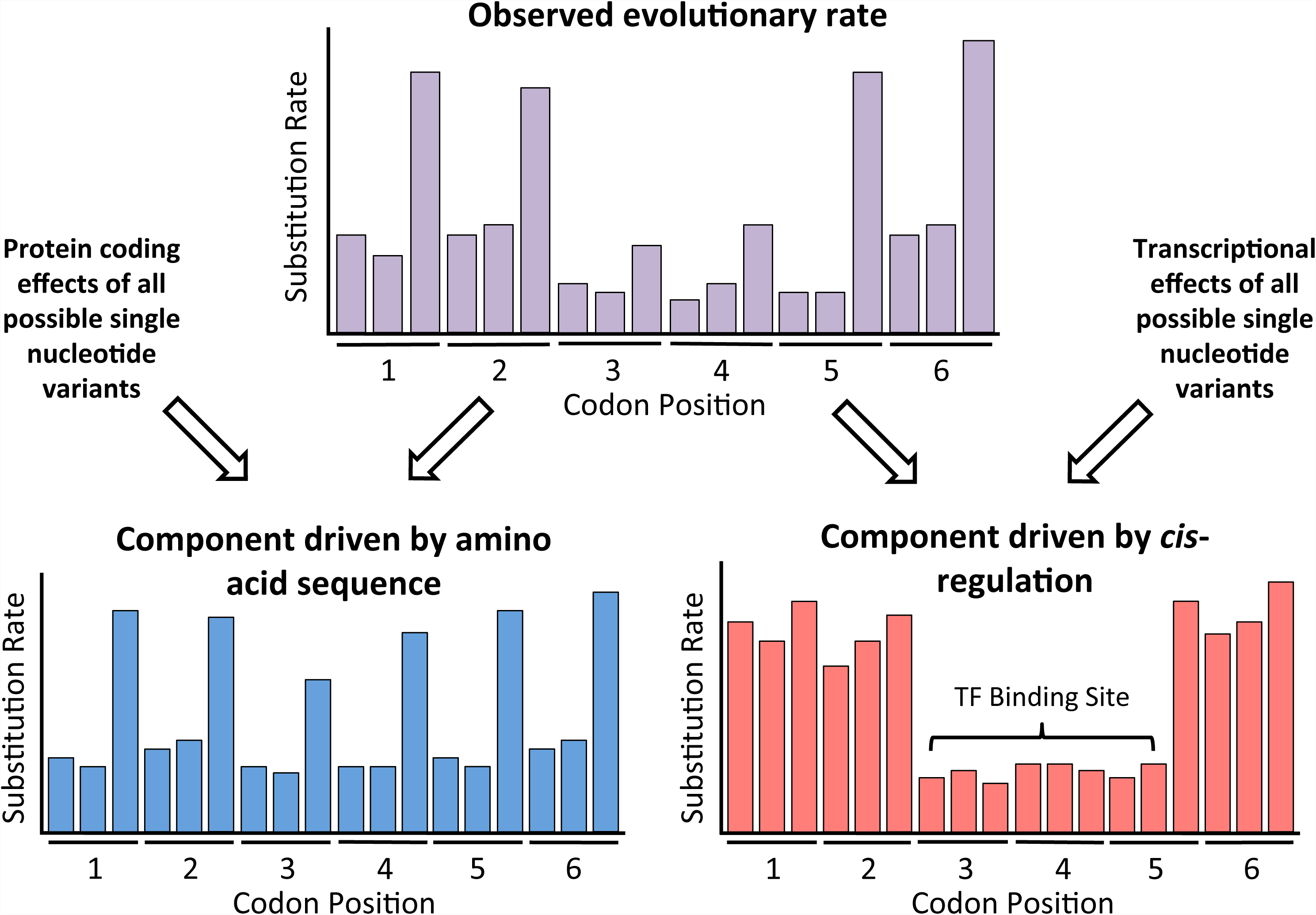
Outline of our approach. For each possible substitution in an exonic enhancer, we determined the evolutionary rate across placental mammals, as inferred from the rate of substitution. We then fit a linear model to determine the relative contribution of two plausible drivers of evolutionary rate: amino acid sequence, where synonymous changes are expected to occur more commonly than nonsynonymous ones; and regulatory impact, wherein positions with high regulatory impact (such as TF binding sites) may dictate lower evolutionary rate.

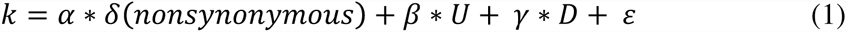

where:

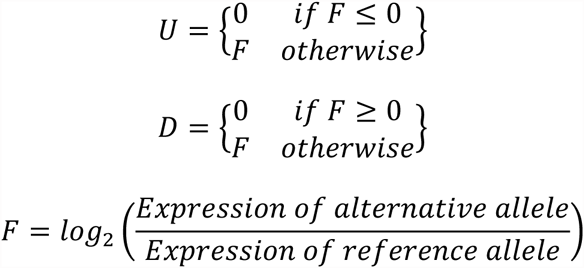

Where *k* is the evolutionary rate of each SNV (measured by rate of that substitution at a given position in the exon); “ nonsynonymous” is a binary indicator of whether each SNV changes an amino acid; “ U” (up-regulation) and “ D” (down-regulation) are the fold-changes in enhancer activity caused by each SNV; and *ε* is an error term. Our goal is to find the coefficients α, *β* and γ that best explain the observed evolutionary rates; negative coefficients would reflect negative selection on each respective type of change, whereas positive coefficients would be expected from a predominance of positive selection. This model is attractive because it naturally accounts for substitutions with effects on both protein sequence and enhancer activity, and it can be extended to more complex models (discussed below). Changes in enhancer activity are split into two separate terms to maximize the linearity of the relationships (with two separate terms, we can capture scenarios such as negative selection on both up- and down-regulation, as well as positive selection for up-regulation coupled with negative selection for down-regulation; these scenarios cannot both be captured with just one regulatory variable in a linear model).

Regulatory effects for each possible SNV were measured in saturation mutagenesis experiments performed by Birnbaum et al. (2014). Their study identified three human exons (*SORL1* exon 17, *TRAF3IP2* exon 2, and *PPARG* exon 6) with strong enhancer activity in liver, and subjected >10^4^ randomly mutated versions of each exon to a massively parallel reporter assay in the livers of live mice, which allowed them to infer each SNV’s precise effect on enhancer activity. These SNVs were also tested in HeLa cells, resulting in different spectra of regulatory effects; we performed our analysis on the data from both cell types.

To determine the evolutionary rate at each position in these exons, we identified and aligned orthologous exon sequences from a wide range of placental mammals (30 - 36 for each exon), and used them to reconstruct the most likely ancestral exon sequence at each node of the phylogeny (Fig. 2a, Supplementary Fig. S1). Each substitution that occurred was identified and scored as synonymous or nonsyonymous. The evolutionary rates of the exons varied, with *PPARG* exon 6 showing the highest and *TRAF3IP2* exon 2 showing the lowest overall conservation (Fig 2b). As expected, there was a strong bias toward synonymous substitutions and transitions over transversions.

**Fig. 2.**
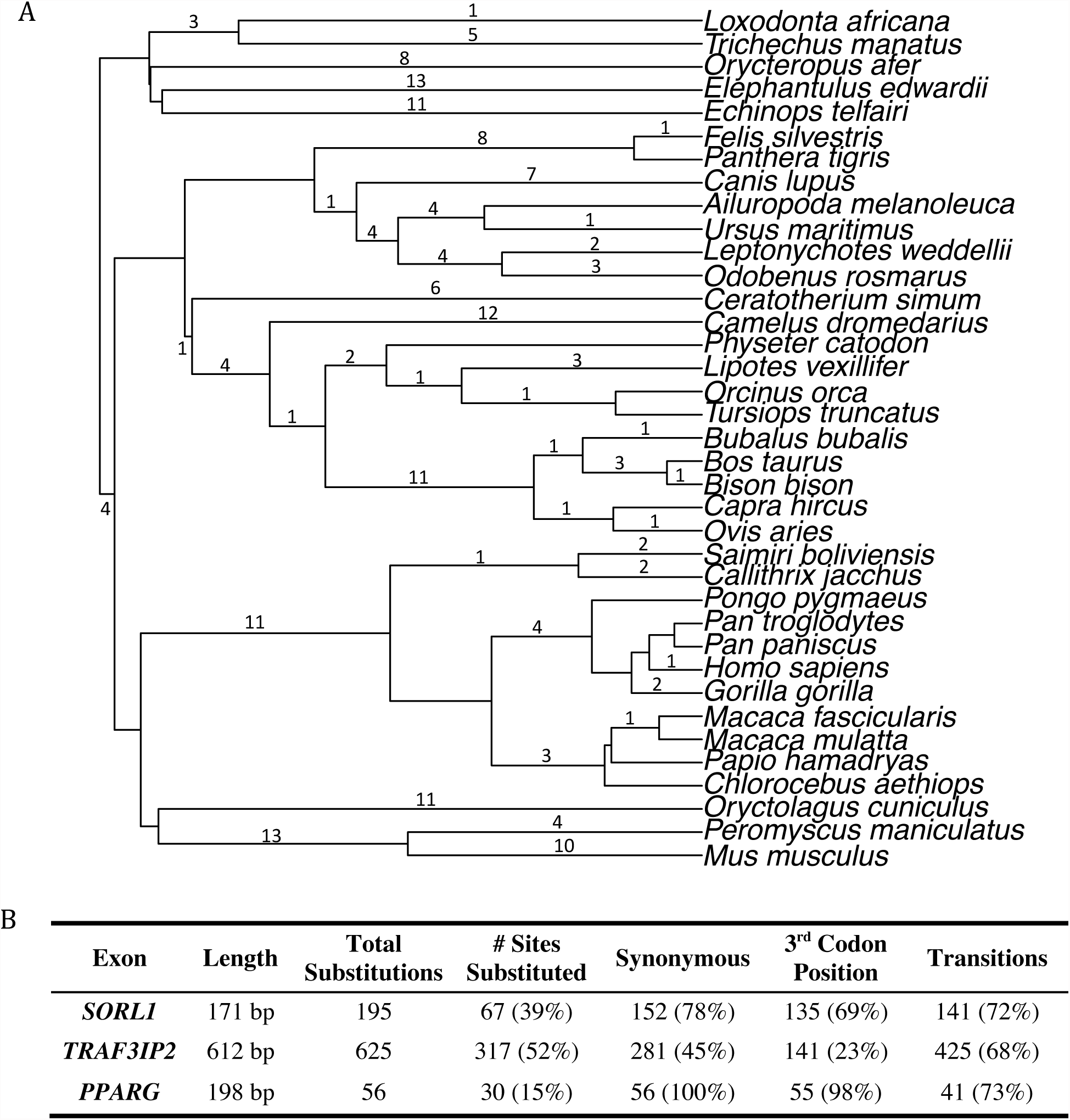
Exonic enhancer evolution. (A) Phylogenetic tree of the mammalian species for which we analyzed the *SORL1* exon 17 sequences. Numbers indicate the number of point mutations identified in each branch. (B) Summary of the substitution patterns for each of the three exons analyzed. Percentages for ‘# Sites Substituted’ are relative to the total number of sites in the exon; percentages for the last three columns are given relative to the total number of substitutions.

In fitting the linear model (1), we determined the three coefficients and their associated p-values for each of the three exons in both liver and HeLa cells (Table 1). For all three exons in both cell types, we found highly significant negative coefficients for the nonsynonymous term (indicating negative selection on protein sequence; all p < 3×10^−11^), but no significance for the regulatory terms (all p > 0.05). This indicates that all detectable selection is acting solely on the protein sequence; when considering the effects of both coding and regulatory changes, the enhancer activities do not contribute significantly to substitution rates.

**Table 1.** Results from linear regression analysis.

| Cell Type | Equation Parameter | *SORL1*<br>Exon 17 |  | *TRAF3IP2*<br>Exon 2 |  | *PPARG*<br>Exon 6 |  |
| --- | --- | --- | --- | --- | --- | --- | --- |
|  |  | Coeff. | P-value | Coeff. | P-value | Coeff. | P-value |
| Liver | Nonsyn. | -0.0167 | 2.2e-13*** | -0.0052 | 2.4e-11*** | -0.0057 | 1.3e-13*** |
|  | Up reg. | -0.0031 | 0.597 | -0.0008 | 0.711 | -0.0026 | 0.232 |
|  | Down reg. | 0.0013 | 0.717 | 0.0010 | 0.476 | -0.0001 | 0.934 |
| HeLa | Nonsyn. | -0.0162 | 1.1e-12*** | -0.0052 | 2.0e-11*** | -0.0056 | 2.5e-13*** |
|  | Up reg. | 0.0275 | 0.051 | 0.0010 | 0.816 | -0.0015 | 0.640 |
|  | Down reg. | 0.0080 | 0.446 | -0.0022 | 0.573 | -0.0005 | 0.791 |

We then explored several additional approaches for fitting the linear model. First, to test for synergistic effects of regulatory and nonsynonymous SNVs, we added pairwise interaction terms to the original model (Table S1). Second, we combined the up- and down-regulation terms from the original model into a single term for regulatory effect (Table S2). Third, we examined the effect of including only up- or only down-regulation, instead of both (Table S3, S4), as well as leaving out the nonsynonymous term (Table S5). Fourth, we replaced our binary nonsynonymous variable, which indicates whether a substitution is synonymous or nonsynonymous, with a variable that further discriminates between conservative vs. radical nonsynonymous changes (Table S6). Finally, because these enhancers have been found to be active in human and mouse (but may not be active in more distantly related mammals) (Birnbaum et al. 2014), we restricted the analysis to just primates and rodents (Table S7). In all cases, these modified linear models confirmed our initial results.

These results suggest that there is little selection on the regulatory functions of these exonic enhancers. This is in stark contrast to the three noncoding enhancers studied by Smith et al (2012), where strong selection was detected on each enhancer. The saturation mutagenesis data used in each study are quite similar (Patwardhan et al. 2012, Birnbaum et al. 2014), having been produced by the same group using the same *in vivo* reporter assay. Therefore the lack of detectable selection on the activities of all three exonic enhancers is unlikely to be explained by an inability of this type of data to reveal selection.

If selection is not acting on these enhancer activities, then a reasonable question is why do these act as such strong enhancers; did they acquire this activity simply by neutral drift? While this may be unlikely to evolve within any given exon, it is important to note that these three exonic enhancers were selected as being among the strongest such elements from a genome-wide screen (Birnbaum et al. 2014). It is not hard to imagine that across the entire human genome, a handful of exons might have evolved enhancer activity simply by chance. However it is also important to note that our approach measures selection across an entire phylogeny; if strong selection was present only in recent human evolution (despite the enhancer being present in both primates and rodents (Birnbaum et al. 2014)), we would not detect it.

Our framework for disentangling sources of selection has a number of limitations. First, it is limited to sequences for which we have saturation mutagenesis data. The number of such studies is small, though rapidly growing (Patwardhan et al. 2009, 2012; Kinney et al. 2010; Melnikov et al. 2012; Kwasnieski et al. 2012; Kheradpour et al. 2013; Birnbaum et al. 2014; Metzger et al. 2015). Second, our approach does not account for epistatic interactions, transacting differences between species, artifacts caused by the plasmid context, or differential effects of SNVs across environments or tissues (see Smith et al. 2012 for further discussion). It is likely that these limitations will be addressed by future saturation mutagenesis studies that examine SNV combinations (c.f. Patwardhan et al. 2012) and additional tissues/environments. In addition, the recent invention of “saturation editing” of the genome now allows SNVs to be studied in their natural chromosomal context (Findlay et al. 2014).

Since our analysis only covers three exonic enhancers, we cannot extrapolate these findings to the rest of the genome. However it is nonetheless striking that even among three of the strongest human exonic enhancers—where one might expect to have the greatest chance of finding selection to maintain their function—the only detectable selection is to conserve amino acid sequence. Therefore we interpret these results as supporting the conclusion of Xing and He (2015), that exonic enhancers likely do not play a major role in protein evolution.

Our framework for disentangling sources of selection is quite flexible, and can be applied to any instance of a sequence encoding distinct functions—such as exonic splice enhancers and silencers, exonic miRNA binding sites, overlapping TF binding sites, etc. As saturation mutagenesis studies become more commonplace, our approach may help distinguish between sequences that only appear to be “functional” because they have some biochemical activity, as opposed to those that are truly important for organismal function and fitness (Graur et al. 2013).

## Methods

We used BLAST (Altschul et al. 1990) to identify orthologs of each exon among placental mammals, using the human sequence as the query. We aligned the sequences with ClustalW2 (McWilliam et al. 2013). The phylogenies were extracted from the best dates mammalian supertree generated by Bininda-Emonds *et al*. (2007). We dropped from the analysis any sequences from mammals not found in this phylogeny, except for those that had a close relative represented in the phylogeny (*Chlorocebus aethiops* for *Chlorocebus sabaeus*, *Pongo pygmaeus* for *Pongo abelii*, *Vicugna vicigna* for *Vicugna pacos*, *Felis sylvestris* for *Felis catus*, and *Papio hamadryas* for *Papio anubis*). We then reconstructed the ancestral sequences at each node of the tree using FastML (Ashkenazy et al. 2012) with the default settings for nucleotide reconstruction. Due to a lack of sensitivity to indels, a small number of manual adjustments were made to the resulting multiple sequence alignment for *TRAF3IP2* exon 2 (three internal node sequences had 6-bp deletions that were incorrectly filled by FastML to match the sequences of their parent nodes).

Within the phylogeny, we located the most likely branch for each substitution by comparing the sequences at each node. Because the saturation mutagenesis data only covered SNVs, the few cases of insertions and deletions were ignored in this analysis. For each possible substitution, we counted how many times it occurred in the phylogeny (most occurred zero times; very few occurred more than once). Because not all possible substitutions had an equal opportunity to occur, we divided these counts by the frequency of the starting base at that position (an “ A” if the change is A → T, etc.) across all sequences in the phylogeny; e.g. a potential substitution whose starting base is present at a particular position in 40 of the sequences in the phylogeny would be divided by 40 to yield a weighted substitution rate. Any potential substitutions whose starting base never appeared in any sequences were excluded from the analysis, since they would never have had the chance to occur.

We then scored each substitution that occurred as synonymous or nonsynonymous. For potential substitutions that never occurred, we scored them based on the human exon sequence. In some rare instances, the same substitution caused a synonymous change in one branch of the phylogeny, and a nonsynonymous change in a different branch. We scored such cases as whichever type they occurred as most often. Instances that were tied for being both synonymous and nonsynonymous (1, 2 and 0 substitutions for *SORL1* exon17, *TRAF3IP2* exon2 and *PPARG* exon 6, respectively) were excluded from the analysis.

We obtained the saturation mutagenesis data from Birnbaum *et al.* (2014) Table S5. For substitutions that did not involve one of the human reference bases, we calculated the effect size by adding the effect sizes (in log-space) of the two human substitutions that would yield the substitution in question. For example, for a T → C substitution at a position that is an A in human, we add the effects of T → A and A → C to get the effect size for this substitution. Regulatory effect sizes were divided into separate variables for up- and down-regulation (up- regulating substitutions were given a value of zero for down-regulation, and vice versa; see equation 1). For the severity scores in our Table S5, we grouped all amino acids based on their size, polarity and charge (Zhang 2000); transitions between these groups were scored as ‘radical’ (given a score of 2) while transitions within these groups were scored as ‘conservative’ (given a score of 1) when considering nonsynonymous substitutions (where synonymous substitutions are given a score of 0). Instances of nonsynonymous changes that were radical in one branch of the phylogeny and conservative in another were handled in the same way as those tied for being synonymous and nonsynonymous, described above. All linear regression analyses were done using the lm() function in the R statistical package (allowing for a flexible intercept), and reported p-values are from the t-test.

## Acknowledgements

We would like to thank J. Smith and the Fraser Lab for helpful discussions and comments on the manuscript. This study was supported by NIH grants T32 GM007790 (supporting RMA) and 1R01GM097171-01A1.

